# Comparative Pathway Integrator: a framework of meta-analytic integration of multiple transcriptomic studies for consensual and differential pathway analysis

**DOI:** 10.1101/444604

**Authors:** Xiangrui Zeng, Zhou Fang, Tianzhou Ma, Chien-Wei Lin, George C. Tseng

## Abstract

**Motivation:** Pathway analysis provides a knowledge-driven approach to interpret differentially expressed genes associated with disease status. Many tools have been developed to analyze a single study. When multiple studies of different conditions are jointly analyzed, novel integrative tools are needed. In addition, pathway redundancy issue introduced by combining public pathway databases hinders knowledge discovery.

**Methods and Results:** We present a meta-analytic integration tool, Comparative Pathway Integrator (CPI), to address these issues using adaptively weighted Fisher’s method to discover consensual and differential enrichment patterns, consensus clustering to reduce pathway redundancy, and a novel text mining algorithm to assist interpretation of the pathway clusters. We applied CPI to jointly analyze six psychiatric disorder transcriptomic studies to demonstrate its effectiveness, and found functions confirmed by previous biological studies as well novel enrichment patterns.

**Availability:** CPI is accessible online: http://tsenglab.biostat.pitt.edu/software.htm.

## 1 Introduction

In a typical transcriptomic study, a set of candidate genes associated with diseases or other outcomes are first identified through differential expression analysis. Then, to gain more insight into the underlying biological mechanism, pathway analysis (a.k.a. gene set analysis) is usually applied to pursue the functional annotation of the candidate biomarker list. The goal behind pathway analysis is to determine whether the detected biomarkers are enriched in pre-defined biological functional domains. These functional domains might come from one of the publicly available databases such as Gene Ontology (GO) [2] and the Kyoto Encyclopedia of Genes and Genomes (KEGG) [12]. Three main categories of pathway analysis methods have been developed in the past decade. The first method called “over-representation analysis” considers biomarkers at a certain differential expressed (DE) evidence cutoff and statistically evaluates the fraction of DE genes in a particular pathway found among the background genes. Without a hard threshold, the second category “functional class scoring” takes the DE evidence scores of all genes in a pathway into account and aggregates them into a single pathway-specific statistics. The third category “pathway topology” further incorporates the information of gene-gene interaction and their cellular location in addition to the pathway database [13].

Many transcriptomic datasets have been generated with the rapid advances of high-throughput genomic technologies in the past decade. Meta-analysis, a set of statistical methods for combining multiple studies of a related hypothesis, has thus become popular. Yet few methods have been developed for the pathway meta-analysis so far. [20] developed two approaches of meta-analysis for pathway enrichment by combining DE evidence at the gene level (MAPE G) or at the pathway level (MAPE P). In real cases, when multiple datasets for a common hypothesis are available under different conditions (e.g., tissues or labs), it might also be interesting to detect pathways enriched consistently under all conditions (consensual pathways) and pathways enriched solely in one condition but not in the others (differential pathways). One naïve way is to identify the enriched pathways in each study individually and manually check whether a certain pathway is enriched in one or multiple studies. To the best of our knowledge, there is currently no available statistical tool that can achieve this goal automatically and systematically.

Another issue emerges with pathway analysis is the pathway redundancy. The large amount of individual pathways identified can hardly infer the underlying biology directly due to this issue. This kind of redundancy typically occurs in a regular pathway analysis since different pathways may include many overlapping genes. Toolkit DAVID [11] resolved this issue by clustering pathways based on a kappa statistics representing the pathway similarity [6]. But the users still need to manually inspect every pathway in a cluster. Furthermore, due to the long and vague pathway description, users can barely make solid conclusions from the results.

In light of this, we proposed a framework of meta-analytical integration of multiple transcriptomic studies for consensual and differential pathway analysis, wrapped in a tool named *Comparative Pathway Integrator (CPI)*. Our tool incorporates 27 databases, including MSigDB [15], GO, KEGG, or user-defined gene set lists, as reference of pathway analysis. In order to identify both commonly and study-specifically enriched pathways, we applied the adaptively weighted Fisher’s method [14], which is originally developed to combine p-values from multiple genomic studies for detecting homogeneous and heterogeneous differentially expressed genes. Clustering analysis based on the overlapping genes among enriched pathways is applied to remove the level of pathway redundancy. Subsequently, we developed a text mining algorithm to automate the annotation of pathway clusters by extracting keywords from pathway descriptions, which also offers more statistically valid summarization compared to leaving user exploring each pathway in a cluster manually and heuristically. Last, CPI visualize the findings and provide users both text and graphical outputs for intuitive while statistical solid presentation and easy interpretation. An R GUI package CPI has been disseminated into *MetaOmics*, an analysis pipeline and browser-based software suite for transcriptomic meta-analysis [16].

## 2 Methods

### 2.1 Workflow of Comparative Pathway Integrator (CPI)

CPI is a comprehensive tool incorporating several widely accepted mature methods as well as some novel algorithms/approaches. It is mainly composed of three steps (Figure 1). The first step is meta-analytic pathway analysis, which includes pathway enrichment analysis and meta analysis. This step partially resembles the work of R package MetaPath [20]. While MetaPath focuses on detecting consensus expressed pathways, CPI will detect both consensual and differentially expressed pathways, providing extra information on how the patterns of pathway enrichment differ across studies. The second step is pathway clustering. This step aims to reduce the redundancy of the pathways from normally hundreds of enriched pathways to a few pathway clusters. The results are more succinct and interpretable. The third step is text mining based on the pathway names and descriptions to find keywords characterizing the overall information of the cluster. A permutation-based statistical test is proformed to assess if a specific biological noun phrases significantly appear more than by chance. Without this step, it would be difficult to identify the representative characteristics for each cluster and users may still need to eyeball all the pathways in results since the clustering of the pathways does not actually reduce the total number of pathways. Finally we have both graphical output, containing pread-sheet output of p-value matrices of pathways and pathway clustering details including gene composition and keywords.

**Figure 1:**
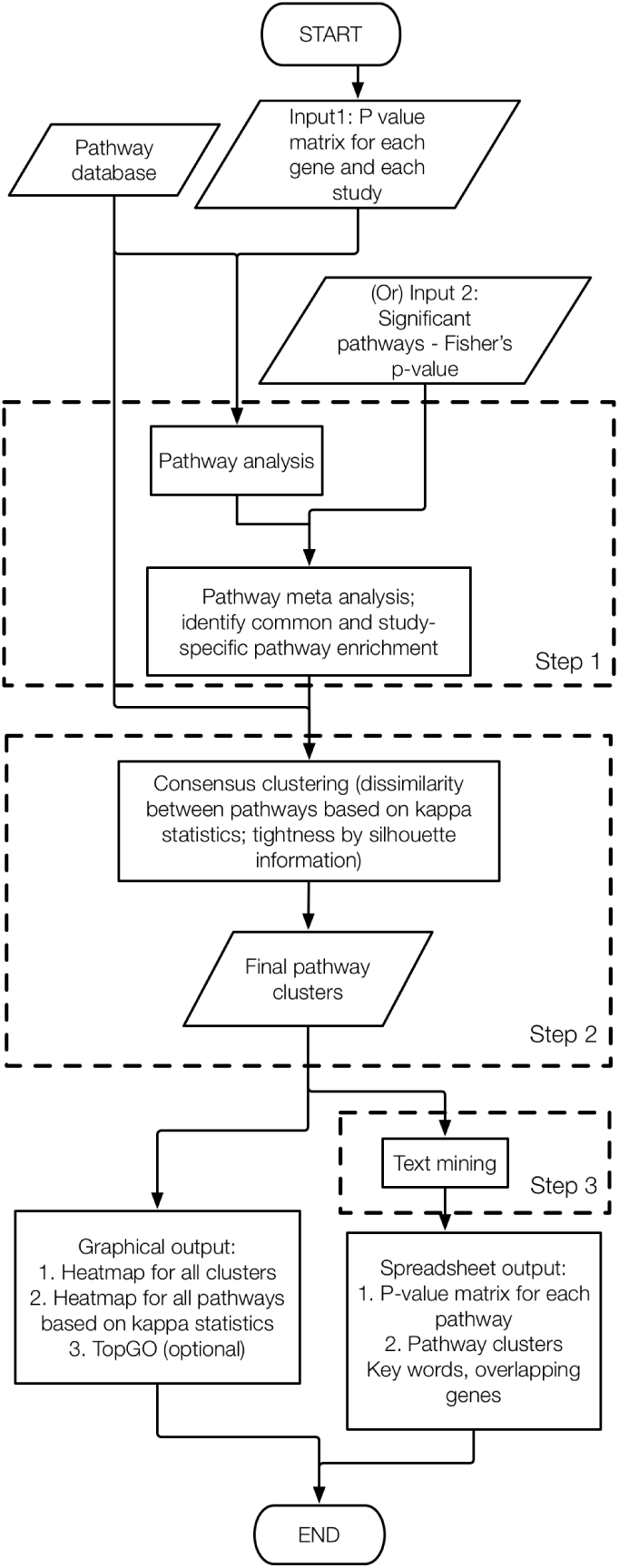
The work flow of CPI

### 2.2 Meta-analytic pathway analysis

Compared with differentially-expressed gene analysis, pathway analysis gives more biological insight to be more systematic and comprehensive. In CPI, we adopt the method of over representation analysis when users input gene list and corresponding p-values from each study. For users who prefer other types of pathway analysis, we also accepts lists of significant pathways and the corresponding p-value as user input.. Given pathway enrichment results, we perform Adaptively-weighted Fisher’s (AW Fisher) method [14] as meta-analysis, to identify pathways significant in one or more studies/conditions. AW Fisher method not only increases the statistical power, but also gives a binary weight for each pathway, indicating the significance of the pathway in each study/condition. Given a user-specified q-value cutoff, we have a list of significant pathways, with some of them commonly significant across studies/conditions while some of them significant in specific studies/conditions.

### 2.3 Pathway clustering for reducing redundancy and enhancing interpretation

Because of the nature of pathways (e.g., hierarchy structure), many genes are shared among different pathways. This redundancy often reduces the interpretability of the result from pathway enrichment analysis. In CPI, we perform pathway clustering method to reduce the redundancy among pathways. The similarity between different pathways is calculated based on Kappa statistics [22], which depends on how many genes are mutual and exclusive among those pathways. The kappa statistics represents the distance between two pathways based on the genes composing each pathway. A distance matrix is then defined based on similarity matrix, composing the distance between each pair of pathways. Consensus clustering [17] is used for clustering purpose. Following the original consensus clustering method, an elbow plot and consensus CDF plot are generated to assist users to decide the number of clusters (Figure 2).

**Figure 2:**
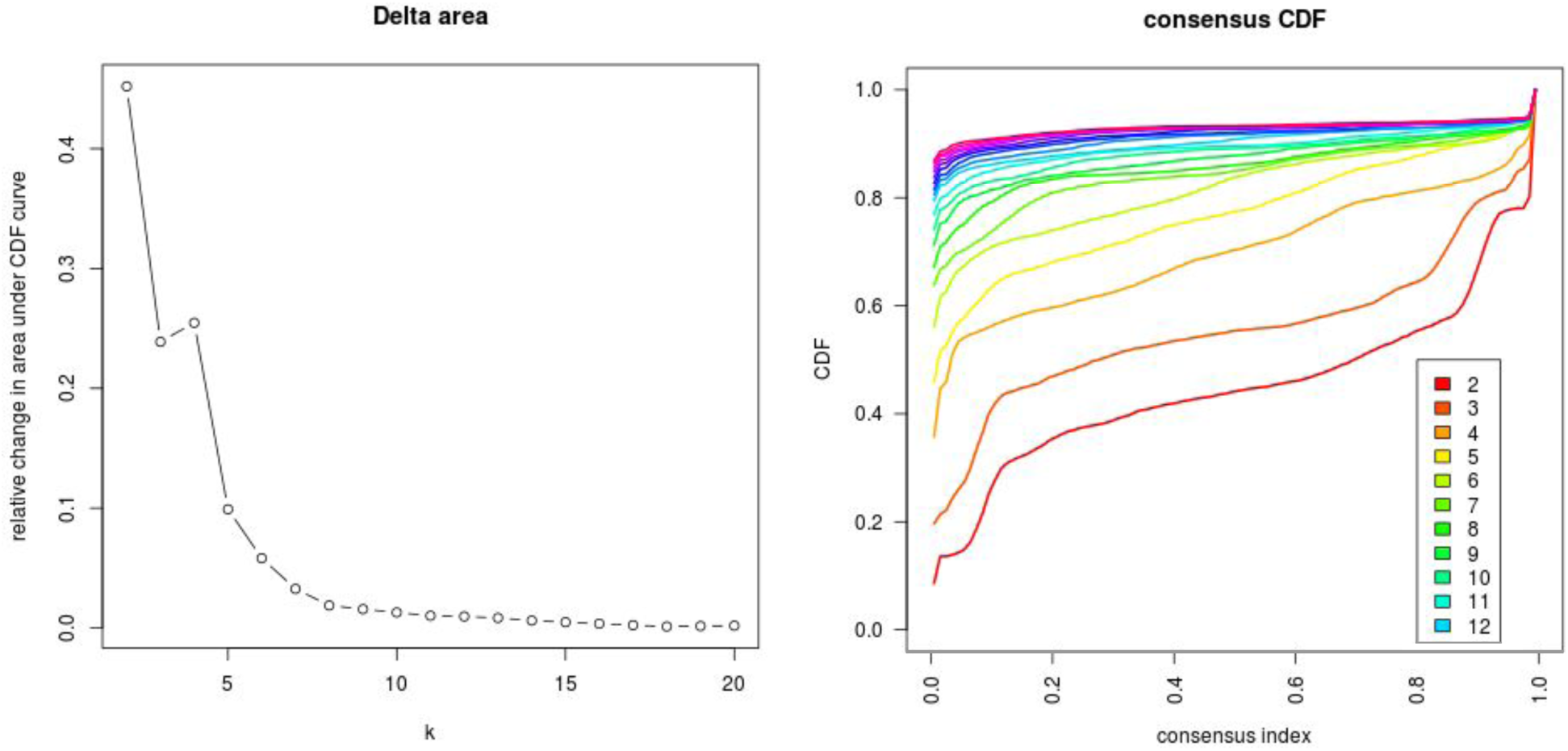
Plots used to assist decision on total cluster number a) elbow plot. b) consensus CDF plot

**Figure 3:**
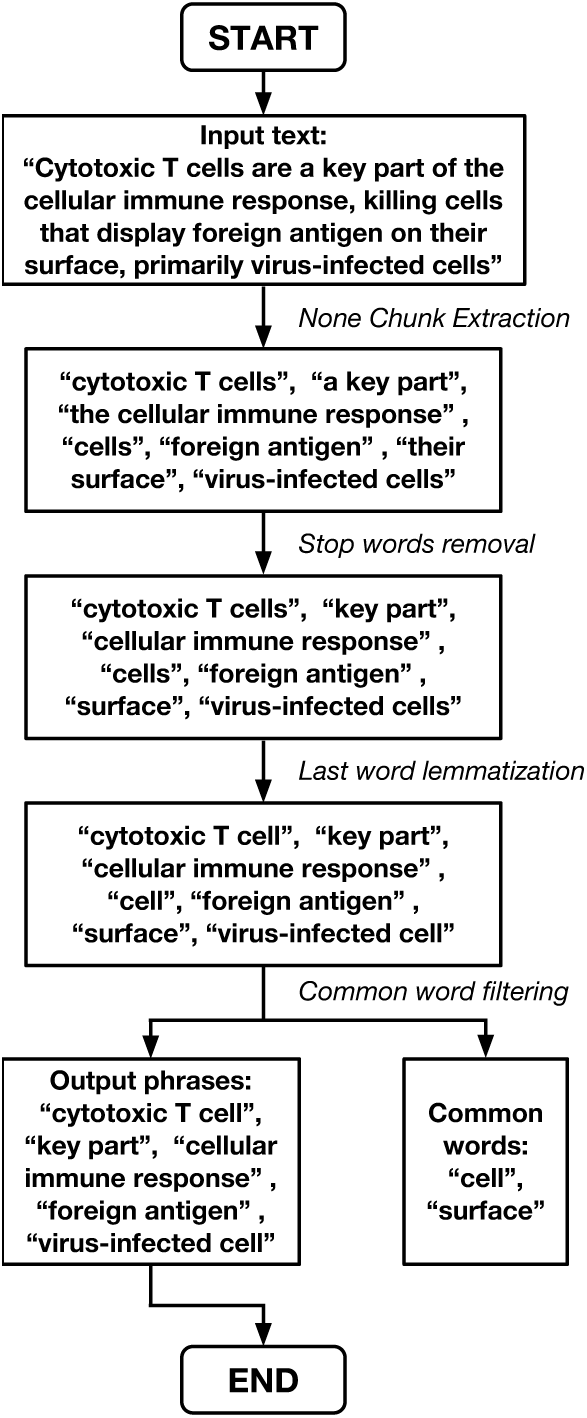
An example of text processing

For most clustering algorithm including the consensus clustering we adopted, each pathway will be forced to cluster into one group of pathways, no matter it’s scattered or not compared to the pathways within the same cluster. We allow scattered pathways to form singletons, when its gene composition is different from representative pathway clusters, to avoid adding noise to other existing clusters. To improve the tightness of the cluster, we further calculated the silhouette width [19], a measure of how tightly each pathway is grouped in its cluster, and removed the scattered pathways with low silhouette width iteratively until all pathways’ silhouette width is above a certain cutoff (we choose 0.1 in our experiments). The removing cutoff for silhouette width is estimated empirically based on its distribution from our multi-disease dataset. For singletons identified, we collected them into another cluster instead of filtering them out because those pathways might be interesting in terms of unique genes composition.

### 2.4 Text mining for automated annotation of pathway clusters

#### 2.4.1 Question description

Reducing the redundancy of pathways by clustering *per se* gives limited summary of the pathways. Since the user usually need to go through most of the pathways in a cluster to grasp an idea and interpretation of the contents in the cluster, the interpretation is not quantitative and largely rely on the biological knowledge of the user. Therefore, we need a more rigorous and statistically meaningful summary of the cluster. The above goal is expressed here as the text mining for key noun phrases for each cluster: which noun phrase appears more frequently in a certain cluster than they usually would? We will therefore treat these noun phrases as the key entities for that cluster. The entity is counted based on the number of pathways containing it, rather than the frequency of it appearing in all pathway descriptions in a cluster. For instance, in a certain cluster, if “T cell” occurs in 3 pathway descriptions (three times in pathway #1, twice in pathway #2 and once in pathway #3),“T cell” is counted 3 times occurrence even if it appears 6 times in total.

#### 2.4.2 Pathway-Phrase matrix

For a pathway description, first, we extracted unique noun phrases from it. This step was done using the noun chunk extraction function from Python package *spaCy* [10]. *spaCy* is an industrial strength text-mining package employing a large library database as well as some machine learning algorithms to detect information from texts. The stop words in English, such as “the”, “a”, “that”, which are very common and carry no important information, are removed from those noun phrases. This step was done using the English stop words database from Python package *NLTK* (Natural Language Toolkit) [4], the Python package with the largest text mining database. After removing all stop words, the last word of each noun phrases, i.e., the central noun of a noun phrase, is lemmatized (converting plural form to singular form) if it is in plural form. This step was done using the lemmatizer function in *NLTK*. The top 5000 common English words are filtered out from the result noun phrases of length one. A text mining process of an example sentence is shown in Figure 4.

**Figure 4:**
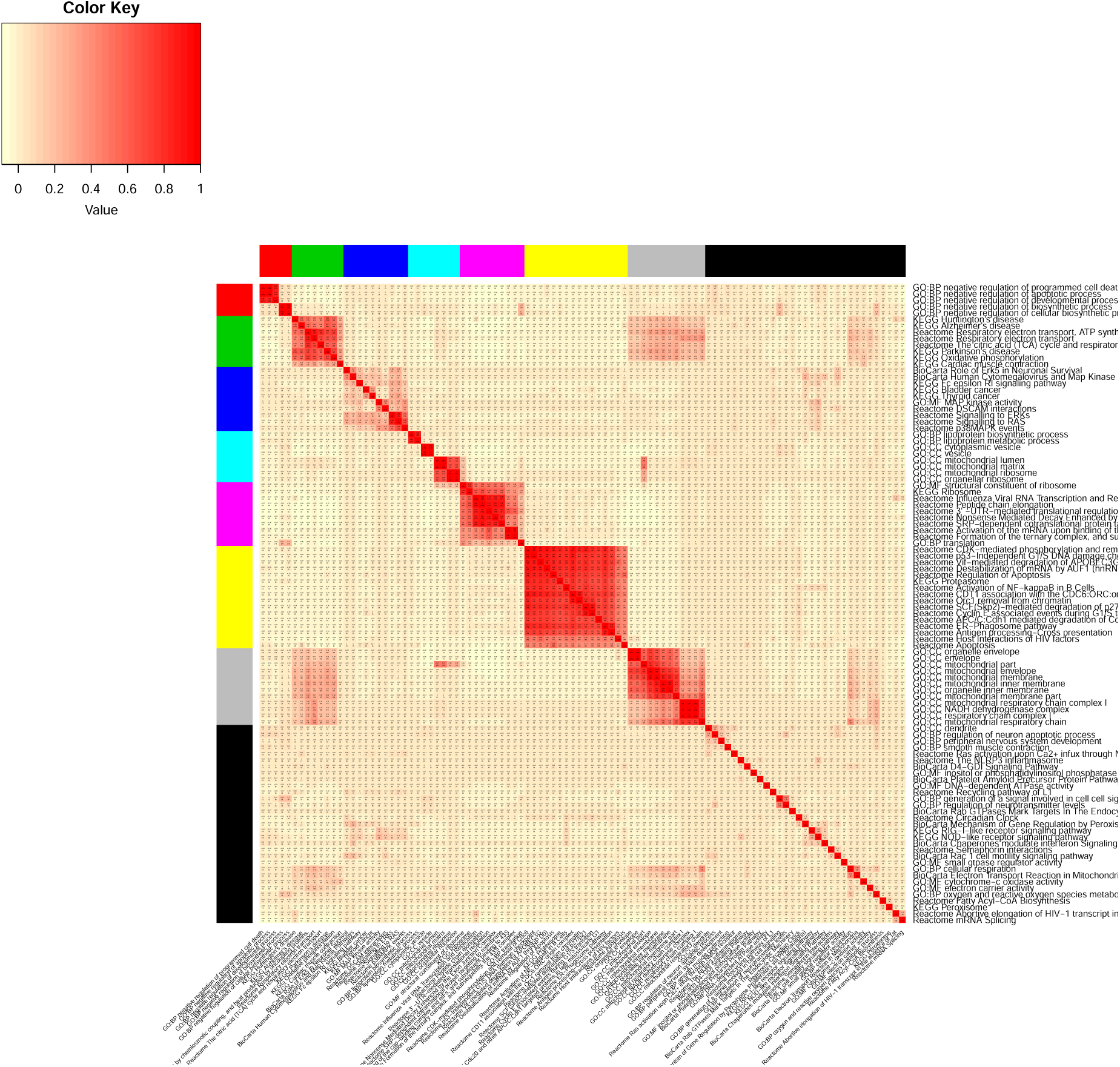
Heatmap of logged p-value of pathways for all clusters using four default databases.

In total, we provide 25 pathway databases (GO, KEGG, BioCarta, Reactome, Phenocarta and etc.) with 26801 pathways. The above formalizing and filtering steps were repeated 26801 times to generate standard form noun phrases for all pathways. Based on the results, we constructed a binary matrix where each row being a noun phrase and each column being a pathway with element *x_ij_* = 1 indicating the pathway description *j* contains the noun phrase *i*.

Once the matrix was constructed, Python package *Vocabulary* was used to identify synonyms from row names of the matrix (noun phrases). When a pair of synonyms are identified, the phrase with lower occurrence in all pathways are combined with the phrase with higher occurrence. Then the row of less occurred phrase was deleted. Since in later cluster text mining, a phrase needs to at least occur in two pathways to be considered, all the rows of phrases which occurred only once in 26801 total pathways are deleted. As a result, a matrix of 24170 rows and 26801 columns was constructed. For later penalized permutation test, the above text mining matrix construction procedure was also applied to pathway names of 26801 pathways. A similar matrix of 24170 rows and 26801 columns *x_ij_* = 1 indicating the pathway term name *j* contains the noun phrase *i*.

#### 2.4.3 Test statistics

A simple strategy to test for the significance of a phrase in a cluster is by simple counting and conduct Fisher exact test. Yet we found this method to be less powerful and biological justifiable from real data analysis, because the phrases in the term name or a shorter description of a pathway are designed to be more representative than those in a full or longer description. We therefore decide to penalize on words found in longer description. We down-weighted the phrase count by assigning a score between 0 and 1 to each pathway *j* to indicate whether it contains phrase *i*:

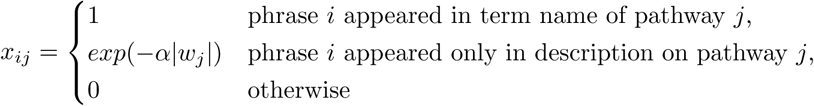

Where *|w_j_|* is the number of unique noun phrases in description of pathway *j*; *α* is a parameter controlling the intensity of penalty. The greater *α* is, the greater the penalty is on longer description. When *α* equals to 0, there is no penalty and our test simplifies to be equivalent to Fisher exact test. And then we define cluster score *T_i_*(*C*) to be the sum of scores of pathways in the cluster, i.e. for phrase i in a pathway cluster *C*, we have our test statistics:

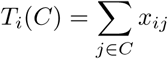

#### 2.4.4 Permutation test

To test for the null hypothesis that a phrase is not enriched in a certain cluster, we adopt a permutation test. The basic argument of constructing the permutation distribution for the test statistics, under the null hypothesis, is that all phrases occur equally frequent across all clusters, including the unpermuted data. So for each phrase *i* in the t*th* permutation, pathways are randomly sampled to form subset *S_t_* with the same cluster size as *C*. Test statistics *T_i_*(*S_t_*) is recomputed at the end of each permutation. The operation is then repeated for a large number of times (say, *T* times). At last, all *T_i_*(*S_t_*) are ranked together with the original data *T_i_*(*C*). And the p-value (later transformed to q-value by BH procedure) could be calculated thereby, indicating how extremely frequent phrase *i* is seen in cluster *C*.

#### 2.4.5 Graphical and Spreadsheet Output

Suppose we already have the clustered pathways and the AW Fisher p-value for each pathway, in this final step we aim to help users understand the overall pattern of pathways significance better, by visualization approaches including heatmaps of p-value for each pathway under each condition, heatmap of kappa statistics for each pathway.

#### 2.4.6 Datasets and databases

We provide 22 homo sapiens pathway database including 14 pathway database from MsigDB, 2 database from Connectivity map, transcription factor target database JASPAR, Protein-Protein interaction database and 3 microRNA databases as options for enrichment analysis.

In addition, Gene Ontology, KEGG database for organism Mus musculus and Saccharomyces cerevisiae, JASPAR database for organism Mus musculus are provided.

## 3 Results

### 3.1 Justifying penalization in text mining

Setting the text mining permutation at 1000 times took a reasonable time for the whole analysis procedure: 9 minutes. Our results also demonstrate the power of penalized permutation test over Fisher’s exact test. Since Fisher’s exact test treats words in description (can be up to 1500 words) the same as words in pathway names (usually less than 15 words), signals of common biological words in pathway names will be masked due to its more frequent occurrence in pathway descriptions, and some non-informative words in descriptions could be detected as keywords falsely. For example, in our text mining results of the two tests, *ATP* in the mitochondrial ATP activity cluster was ranked low in the Fisher’s exact test (r=17) while prioritized in permutation test (r=1). Some meaningless words such as *buolo* and *engelhardt* in cluster 3 was ranked high in Fisher’s exact test (r=5), but ranked low by penalized permutation test (r=37 and r=18). To conclude, for detecting ordinary keywords, penalized permutation test performs roughly the same as Fisher’s exact test. However, for detecting keywords frequently occur in biological vocabulary and filtering out strange jargons penalized permutation test is indeed more powerful than Fisher’s exact test.

### 3.2 Results for real data

To demonstrate its utility, we applied CPI using the default databases (Gene Ontology, KEGG, Reactome and BioCarta) to analyze the transcriptome datasets of three psychiatric diseases of two prefrontal cortex layers. These datasets, provided by Dr. David A. Lewis’ group, was used previously to compare post-mortem tissue dorsolateral prefrontal cortex (DLPFC) layer 3 and layer 5 pyramidal cells’ gene expression level of bipolar disorder, major depressive and schizophrenia patients with matched healthy control [18].

By inputting the top 400 differentially expressed genes in each of the six datasets and default pathways containing 10 to 200 genes, top 100 major pathways of the total 2187 pathways were identified and clustered to 7 clusters with 31 pathways as singleton terms. Of seven clusters, three important clusters with insightful biological meaning are discussed below. The elbow plot and consensus CDF plot (Figure 2) indicate that a cluster number of 7 is reasonable. Because when the number of clusters is greater than 7, the relative change in area under CDF curve of elbow plot slows down (Figure 2a), and the consensus CDF curve flattens out (Figure 2b).

Based on the heatmap output shown in Figure 4 and 5, we found that pathways in cluster 1 are significantly altered in schizophrenia DLPFC layer 5. Cluster 2, 5 and 7 shows similar pattern, and are significantly enriched consensually in both schizophrenia DLPFC layer 3 and 5. Cluster 3 is enriched in the DLPFC layer 5 of two diseases: significantly in major depression disorder, and marginally, bipolar disease. Cluster 4 is solely enriched in the layer 3 of schizophrenia. Cluster 6 is enriched significantly in layer 3 of schizophrenia, and slightly in the layer 3 of schizophrenia and bipolar disease. And we also observed that the singletons on the bottom of the heatmap display no consensual nor differential pattern, as they are scattered pathways.

**Figure 5:**
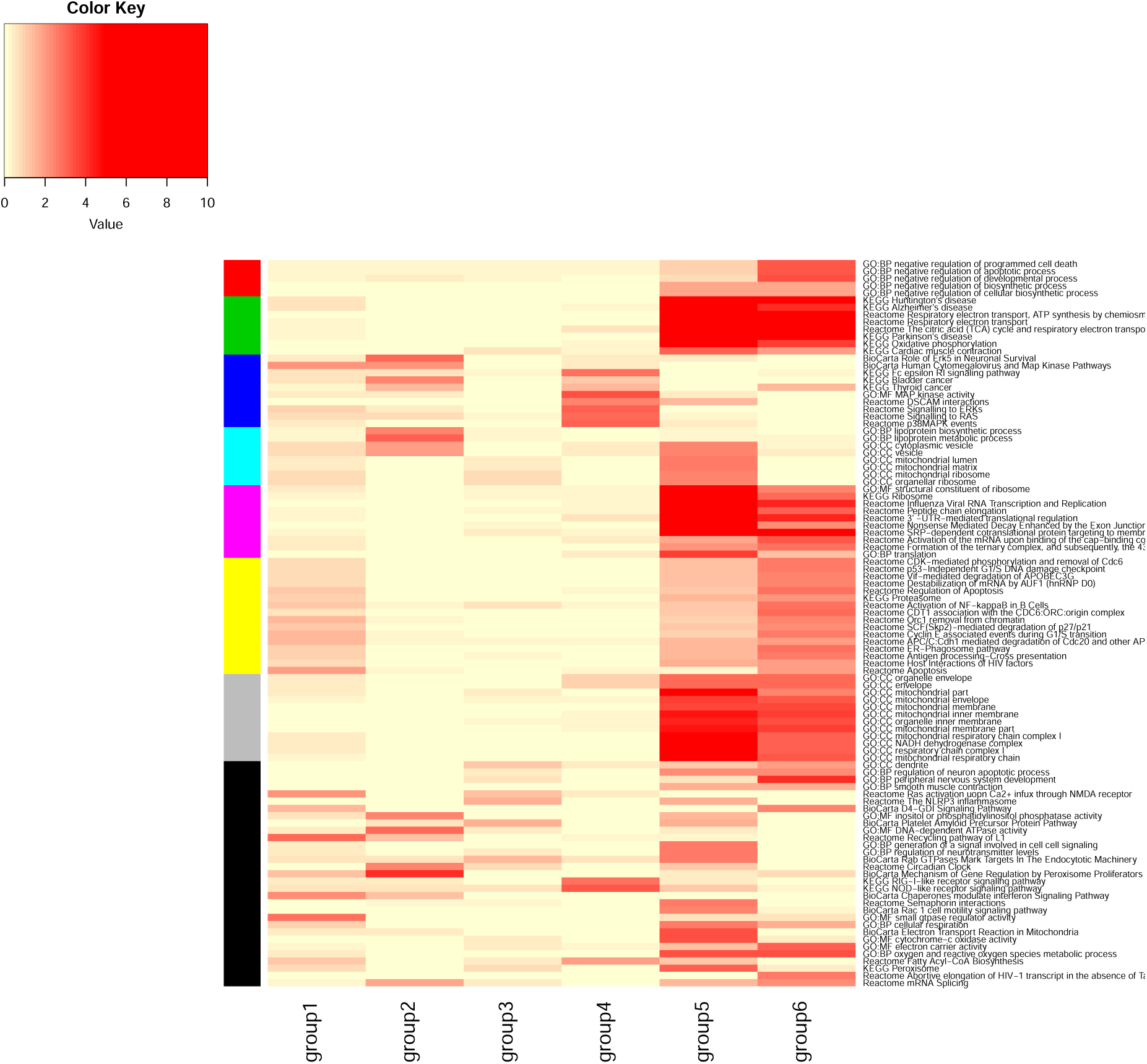
Heatmap of logged p-value of pathways for all clusters using default databases.

To further demonstrate the consensual and differential patterns of our results, we inspect the contents of each cluster with the help of text mining. The information provided from these pathways are diverse, even within a cluster. But those information could be condensed statistically into keywords, with our penalized text mining method.

Based on pathway clustering and text mining results in the clustering summary csv file from step 3, we found that cluster 5 of 8 pathways with keywords under FDR 0.01: *innate immune response, cytokine, MAPK* is significantly altered in major depression DLPFC layer 5 (group 4). Reduced MAPK/ERK signaling in hippocampal area is associated with depressive behavior [7, 9, 8]. It is unknown whether major depressive disorder is associated with altered MAPK/ERK signaling in dorsolateral prefrontal cortex. Our study demonstrated altered MAPK/ERK signaling in major depressive disorder DLPFC layer 5 but not in layer 3 or the other two diseases. It remains an open question how this alteration relates to major depression disorder symptoms. Cluster 6 of 8 pathways with keywords under FDR 0.02: *inner mitochondrial membrane, NADH, ATP synthesis* is significantly altered in layer 3 of schizophrenia DLPFC (group 5) and to a less extent in layer 5 of schizophrenia DLPFC (group 6). This consistent with a previous transcriptomic study of schizophrenia and schizoaffective disorder in Dr. Lewis’ lab [1]. And our results further indicated that the difference in the degree of mitochondrial dysfunction across DLPFC layer 3 and layer 5 was mainly due to different degrees of mitochondrial ATP dysfunction rather than other aspects of mitochondrial dysfunction. Cluster 7 of 33 pathways with keywords under FDR 0.01: *DNA replication, mitosis, ubiquitin* is significantly altered in schizophrenia DLPFC layer 3 (group 5) and to a less extent in bipolar disorder DLPFC layer 3 (group 1) and schizophrenia DLPFC layer 5 (group 6). This validates another previous finding from Dr. Lewis’s lab which associated ubiquitin-proteasome related genes to schizophrenia layer 5 [1]. Furthermore, consensual results across different diseases and layers are shown by the significant alteration of ubiquitin-proteasome system and cell cycle not only in schizophrenia DLPFC layer 5, but also in schizophrenia and bipolar disorder DLPFC layer 3. Similar result from a blood based microarray investigation evidenced preliminarily ubiquitin-proteasome dysregulation in both schizophrenia and bipolar disorder [5]. However, the DE genes contributing to ubiquitin-proteasome dysregulation are different in bipolar disorder and schizophrenia, which may need further biological investigation and interpretation.

## 4 Conclusion

In this article, we explored the approaches for comparative meta-analytic pathway analysis, and developed an integrative platform for this purpose called “CPI”. CPI reduces pathway redundancy to condense knowledge discovered from the results and also conducts text mining to provide statistically solid suggestions on interpreting results. CPI has three major steps. In the first step, users may input either p-value matrix of either genes or pathways for each study to start with. If p-values of genes are inputted, pathway analysis is applied first. Then user may select a q-value cutoff for significant pathways [3]. And those pathways are passed down to meta analysis where AW Fisher is applied to discover consensually and differentially enriched pathways. We suggest 0.2 as our default cutoff, and the users may also choose their own cutoff according to their budget on the follow-up experiments and analysis. In the second step, the pathways are clustered using consensus clustering, with consensus CDF plot and elbow plot assisting users to choose the number of clusters [17]. By default, a group of six clusters were assessed. Silhouette information is used to achieve cluster tightness by moving scattered pathways to singleton. The cutoff of 0.1 is selected using silhouette information [19] distribution from our multi-disease dataset, and results in a reasonable number of singletons (31 out of 100 pathways). In the third step, text mining, users may choose number of permutation iterations based on user’s computation capacity and data size. The computational complexities of the CPI R standalone and GUI version are similar. Using one thread on iX Windows system, with the input of 6 gene lists with total of 84 subjects and 44k genes and the default pathway reference database, setting permutation test iteration to be 1000, one run of CPI takes 9 minutes. Moreover, both versions can be executed in parallel to further speed up the analysis.

CPI has three advantages as compared to previous methods addressing pathway meta-analysis. First, CPI explores consensual and differential expression pattern spontaneously in integrated pathway analysis. Second, CPI clusters pathways by the gene composition to reduce pathway redundancy. Third, CPI uses a statistically valid text mining method to interpret pathway analysis results. In addition, the penalized text mining algorithm by permutation test has shown the advantage over the standard test like Fisher’s exact test based on real data analysis. We applied the tool to multiple psychiatric disorders transcriptomic data. The result identifies multiple pathway enrichment patterns relevant to previously confirmed as well as novel biological functions, such as mitochondrial ATP dysfunction in schizophrenia DLPFC layer 3, ubiquitin-proteasome system dysregulation in schizophrenia and bipolar disorder DLPFC layer 5 and altered MAPK/ERK signaling chain in major depression disorder DLPFC layer 5 [1].

CPI has several limitations. First, for single study pathway analysis, our tool only allows user to apply Fisher’s exact test and KS test, but it can be readily extend to include other methods such as GSEA [21]. Second, our text mining algorithm rely on the descriptions provided by pathway databases. So for those pathway databases that do not provide descriptions, e.g. some pathways in KEGG, text mining algorithm loses advantages. Thirdly, computation time is not ignorable, especially in text mining step. For the real data we applied, each 1000 permutation iterations increase 5 minutes computation time.

In summary, CPI is a meta-analytic tool for discovering commonly expressed and study specific pattern in transcriptomic studies, that will also reduce pathway redundancy and conduct text mining to increase interpretability of the results. CPI is implemented in R, as well as R Shiny, an R based graphical user interface. The R Shiny version is disseminated in MetaOmics can be easily handled by users without programming knowledge [16].

## Funding

This work has been supported by the NIH R01CA190766 and R21LM012752.

